# Impact of temperature on the affinity of SARS-CoV-2 Spike for ACE2

**DOI:** 10.1101/2021.07.09.451812

**Authors:** Jérémie Prévost, Jonathan Richard, Romain Gasser, Shilei Ding, Clément Fage, Sai Priya Anand, Damien Adam, Natasha Gupta Vergara, Alexandra Tauzin, Mehdi Benlarbi, Shang Yu Gong, Guillaume Goyette, Anik Privé, Sandrine Moreira, Hugues Charest, Michel Roger, Walther Mothes, Marzena Pazgier, Emmanuelle Brochiero, Guy Boivin, Cameron F. Abrams, Arne Schön, Andrés Finzi

## Abstract

The seasonal nature in the outbreaks of respiratory viral infections with increased transmission during low temperatures has been well established. The current COVID-19 pandemic makes no exception, and temperature has been suggested to play a role on the viability and transmissibility of SARS-CoV-2. The receptor binding domain (RBD) of the Spike glycoprotein binds to the angiotensin-converting enzyme 2 (ACE2) to initiate viral fusion. Studying the effect of temperature on the receptor-Spike interaction, we observed a significant and stepwise increase in RBD-ACE2 affinity at low temperatures, resulting in slower dissociation kinetics. This translated into enhanced interaction of the full Spike to ACE2 receptor and higher viral attachment at low temperatures. Interestingly, the RBD N501Y mutation, present in emerging variants of concern (VOCs) that are fueling the pandemic worldwide, bypassed this requirement. This data suggests that the acquisition of N501Y reflects an adaptation to warmer climates, a hypothesis that remains to be tested.

## INTRODUCTION

The etiological agent of the COVID-19 pandemic is the SARS-CoV-2 virus (1). While it will likely take years to understand the spread of SARS-CoV-2 infection in the human population, several factors could be modulating transmission dynamics and are currently being heavily scrutinized. As for other respiratory viruses, host immunity, population density, human behavioral factors, humidity and temperature likely modulate its transmission (2–7). Different steps in the replication cycle of coronaviruses could be affected by such factors, particularly viral entry. This process is mediated by the viral Spike (S) glycoprotein. The Spike glycoprotein uses its receptor binding domain (RBD) to interact with its host receptor angiotensin-converting enzyme 2 (ACE2) (8–10). Cleavage by cell surface proteases or endosomal cathepsins (8,11,12) releases the fusion peptide and triggers irreversible conformational changes in the Spike glycoprotein enabling membrane fusion and viral entry (13, 14).

Airway transmission of SARS-CoV-2 is confronted to the temperature gradient that exists in human airways. In the nasal mucosa it is around 30 to 32°C, it moves up to 32°C in the upper trachea, and around 36°C in the bronchi (15, 16). Emerging results strongly suggest that the Spike glycoprotein of SARS-CoV-2 evolved to replicate in the upper airways (17). This was linked to Spike stability which was enhanced by the introduction of the D614G mutation (18–20) but also enhanced its use of cell-surface and endosomal proteases (17, 21). Additionally, recent studies have noticed an increase replication of SARS-CoV-2 in primary human airway epithelial cells at 33°C compared to 37°C (22), while higher temperatures (39°C-40°C) decreased overall viral replication (23). Since it was previously documented that affinity of viral envelope glycoproteins for their receptor is modulated by temperature (24), herein, we evaluate to what extent temperature affects the interaction of SARS-CoV-2 Spike with ACE2, and concomitantly, its effect on viral attachment.

## RESULTS

### Conservation of ACE2-interacting residues among SARS-CoV-2 Spike isolates

We first evaluated the conservation of the ACE2-binding site on the SARS-CoV-2 Spike. Based on previously published structural data (25), ACE2 interacts with 17 key residues on the RBD primarily located on the receptor-binding motif (RBM). To determine the degree of conservation of these residues, we used the COVID-19 CoV Genetics program (26) and analyzed SARS-CoV-2 sequences deposited in 2019-2020 or in 2021 (up to June 18^th^ 2021). Over the 2019-2020 period, no major sequence variations were observed except for the N501Y mutation (4.4%), which started to arise at the end of the year in at least three independent lineages of interest (B.1.1.7, B.1.351, P.1) (27–29). In 2021 sequences, most residues were still found to be >99% conserved except for variations found at residues 417 (K417N, 1.5%; K417T, 2.6%) and 501 (N501Y, 65.2%), with the latter becoming predominant among all the deposited sequences in 2021 (Fig 1A). These mutations are found in emergent variants of concern (VOCs), including the B.1.1.7 (N501Y), B.1.351 (K417N/N501Y) and P.1 lineages (K417T/N501Y) (30). Among them, the B.1.1.7 lineage (also known as alpha variant), which was first identified in the United Kingdom was shown to have increased infectivity and transmissibility (31, 32). This lineage spread rapidly and is the major circulating strain in early 2021 worldwide, replacing the D614G strain which was predominant in 2019-2020 (Fig 1B) (33). Sequence variations were also found in other RBM residues (Fig S1), notably mutations L452R (8.8%), E484K (7.7%), T478K (5.9%), S477N (2.2%) and N439K (1.2%) which are also found in other various VOCs (34–38) and were shown to either increase infectivity or promote the evasion of antibody responses (30,34,38–42).

**Figure 1.**
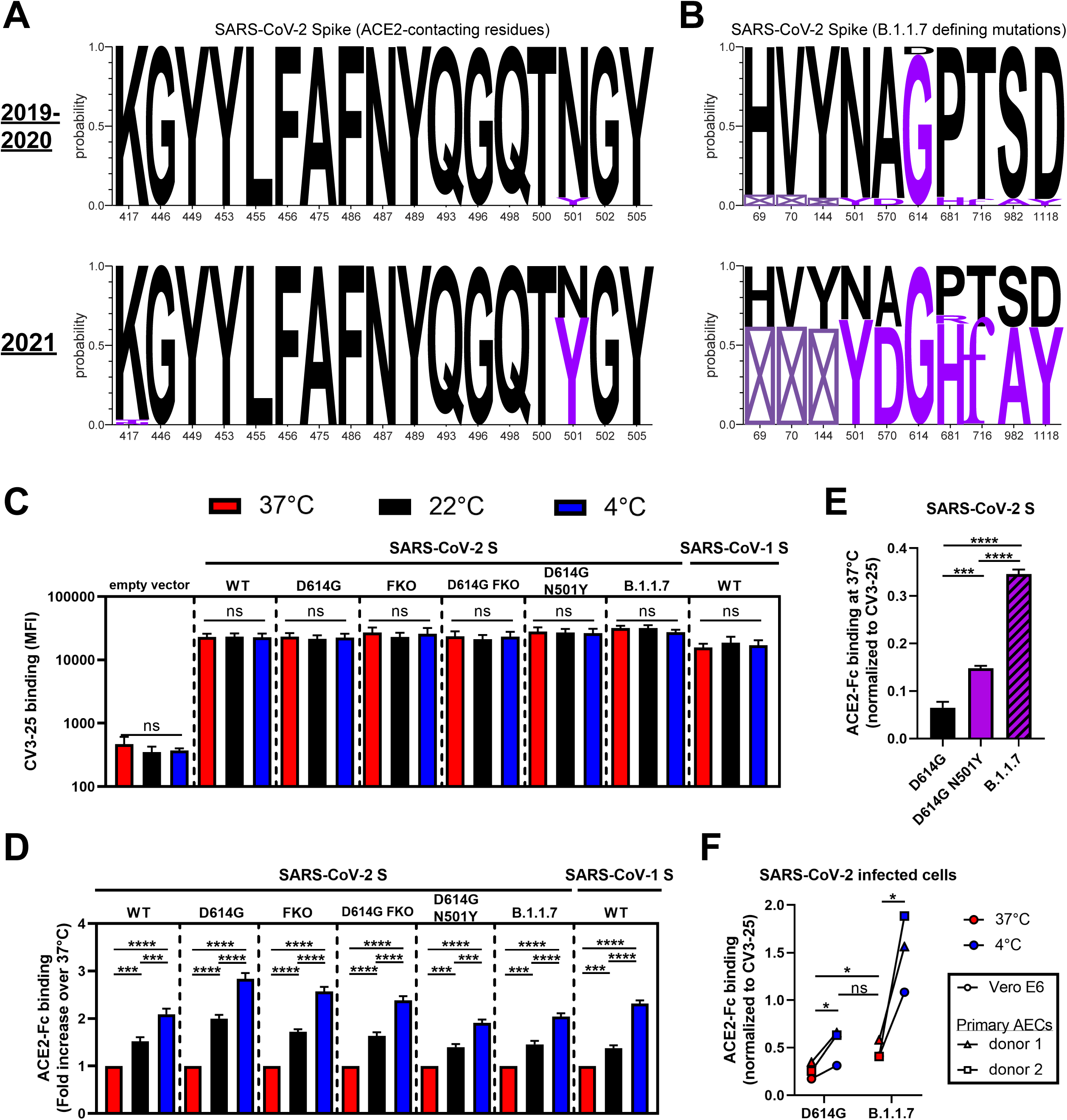
Enhanced binding of ACE2 to SARS-CoV-2 Spike at low temperatures. (**A-B**) Logo depictions of the frequency of SARS-CoV-2 Spike residues known to (**A**) contact with ACE2 or (**B**) corresponding to B.1.1.7 defining mutations. Worldwide sequences deposited in the GISAID database in 2019-2020 and in 2021 (up to June 18^th^, 2021) were aligned using the COVID CoV Genetics program. The height of the letter indicates its frequency over total sequences. Residues corresponding to the WIV04 reference sequence are shown in black and residues corresponding to VOCs are shown in violet. A box with a cross inside (☒) indicates the presence of a residue deletion. (**C-E**) Cell-surface staining of transfected 293T cells expressing SARS-CoV-2 Spike (WT, D614G, Furin KO, D614G Furin KO, D614G N501Y or B.1.1.7 variant) or SARS-CoV-1 Spike (WT) using (**C**) CV3-25 mAb or (**D-E**) ACE2-Fc. (**F**) Cell-surface staining of Vero E6 or primary human AECs from two healthy donors infected with authentic SARS-CoV-2 viruses (D614G or B.1.1.7 variant) using ACE2-Fc. (**C,E,F**) The graphs shown represent the binding of primary antibodies performed at 37°C, 22°C or 4°C. Panel **D** is depicting the binding of ACE2-Fc at 37°C only. ACE2-Fc binding was normalized to CV3-25 binding in each experiment at each temperature. The graphs shown represent the median fluorescence intensities (MFI). Error bars indicate means ± SEM. These results were obtained in at least 3 independent experiments. Statistical significance was tested using (**C-E**) one-way ANOVA with a Holm-Sidak post-test or (**F**) a paired T test (*p < 0.05; ***p < 0.001; ****p < 0.0001; ns, non significant).

### Low temperatures increase SARS-CoV-2 Spike-ACE2 interaction

To measure the effect of temperature on the Spike-ACE2 interaction, we use a system where we express the full-length native Spike at the surface of cells and measure its interaction with the ACE2 receptor using a recombinant ACE2-Fc chimeric protein. This recombinant protein is composed of an ACE2 ectodomain linked to a human IgG1 Fc (43). 293T cells were transfected with a plasmid encoding the SARS-CoV-2 wild-type (WT) Spike (Wuhan-Hu-1 strain). Forty-eight hours post-transfection, cells were incubated at different temperatures (37°C, 22°C and 4°C) before measuring ACE2-Fc binding by flow cytometry. To ensure that any differential recognition was not linked to a temperature-dependent variation in Spike levels, we used the conformational-independent S2-targeting monoclonal antibody (mAb) CV3-25 as an experimental control (44, 45). As shown in Fig 1C, temperature did not alter CV3-25 recognition, indicating that temperature does not affect the overall amount of Spike at the surface of these cells. Therefore, the CV3-25 mAb was used to normalize Spike expression levels among the different mutants or variants (Fig 1D-F).

Interestingly, we observed a gradual increase in ACE2-Fc recognition concomitant with the temperature decrease (Fig 1D), suggesting a temperature-dependent interaction between Spike and ACE2. Since temperature was suggested to also affect Spike stability (17,46,47) which in turn could explain its decreased receptor binding at 37°C, we introduced the D614G change, known to increase trimer stability (17, 19) in combination or not with furin cleavage site mutations (FKO), known to prevent Spike proteolytic cleavage (48). The same stepwise increase in ACE2-Fc binding at lower temperatures was observed with all Spike constructs (Fig 1D), indicating that low temperature can enhance ACE2-Fc binding independently of the strength of association between the S1 and S2 subunits. To extend these results to the Spike of emergent circulating strains, we evaluated ACE2-Fc binding to the Spike N501Y mutant and the Spike from the B.1.1.7 lineage. The N501Y mutation is located at the RBD-ACE2 interface and has been previously shown to strengthen the interaction with ACE2 by inserting an aromatic ring into a cavity at the binding interface (47, 48). Despite significantly higher binding of ACE2-Fc at 37°C (Fig 1E), a similar enhancement was observed with both the N501Y mutant and the B.1.1.7 variant at low temperatures (Fig 1D). To evaluate whether this phenotype was conserved among other Betacoronaviruses, we also performed the same experiment using the closely related SARS-CoV-1 Spike and similar changes were observed (Fig 1D).

To confirm our observations in a more physiological model, we infected a highly permissive cell line (Vero E6) and primary airway epithelial cells (AECs) from two different healthy donors using authentic SARS-CoV-2 virus isolated from patients infected with SARS-CoV-2 D614G or B.1.1.7 (Fig 1F). Using flow cytometry, we discriminated the infected cells using an anti-nucleocapsid mAb and measured the binding of ACE2-Fc at the cell surface. In agreement with results from transfected cells, the binding of ACE2 to cell surface Spike was higher at cold temperature (4°C) compared to 37°C for both D614G- and B.1.1.7-infected cells (Fig 1F). Importantly, ACE2 bound efficiently to the B.1.1.7 Spike at 37°C (Fig 1F). Similar level of binding could only be achieved for the D614G Spike by decreasing the temperature to 4°C (Fig 1F). Overall, low temperatures appear to promote Spike-ACE2 interaction independently of Spike trimer stability and emerging mutations, although the B.1.1.7 variant exhibited a pronounced improvement in binding at warmer temperatures.

### Low temperatures improve the viral attachment of SARS-CoV-2 virions

Next, we investigated the effect of enhanced ACE2 binding at low temperatures on SARS-CoV-2 Spike functional properties, including its ability to mediate viral attachment and fusion, and the subsequent consequences on early viral replication kinetics. To assess viral attachment, we adapted a previously described virus capture assay (49) where we generate lentiviral particles bearing SARS-CoV-2 Spike and look at their ability to interact with ACE2-Fc immobilized on ELISA plates. In agreement with a better affinity for ACE2 at lower temperatures, more SARS-CoV-2 D614G pseudoviral particles were captured at 4°C compared to 37°C (Fig 2A). In line with these results, we also observed enhanced infectivity and cell-to-cell fusion mediated by SARS-CoV-2 Spike D614G at 4°C compared to 37°C, while a marginal increase was seen with an unrelated viral glycoprotein (VSV-G) (Fig 2B-C). Similarly, the capacity of soluble ACE2 (sACE2) to neutralize pseudovirions bearing SARS-CoV-2 Spike D614G was significantly improved when pre-incubating the virus with sACE2 at 4°C when compared to 37°C prior infection of 293T-ACE2 target cells (Fig 2D-E). Similar effects of temperature on Spike-mediated attachment and fusion, and on sensitivity to sACE2 neutralization were observed when using the Spike N501Y mutant or B.1.1.7 variant (Fig 2A-E). To analyze the impact of temperature on viral replication in a more physiological model, we used authentic SARS-CoV-2 D614G viruses to infect reconstituted primary human airway epithelia (MucilAir). Infections were performed at 4°C or 37°C for 30 minutes, virus-containing medium was then discarded to remove any unbound virus before keeping the cells at 37°C for four days. While no significant differences in viral titers were observed at 24h post-infection, viral replication at 96h post-infection was found to be significantly higher when the initial infection was performed at 4°C versus 37°C (Fig 2F). Altogether, this suggests that lower temperatures improve the initial attachment of SARS-CoV-2, which in turn can alter the subsequent kinetics of viral replication.

**Figure 2.**
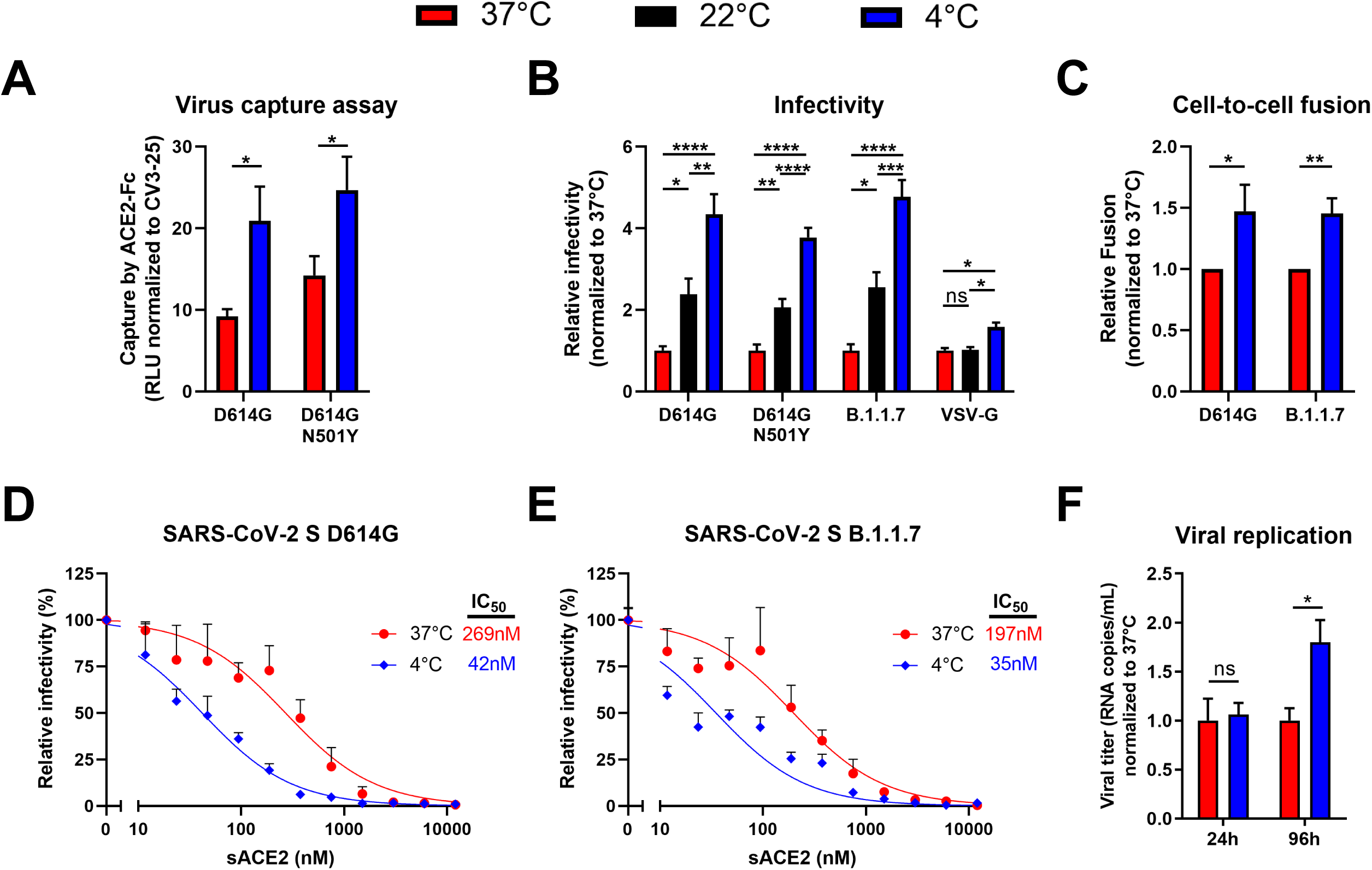
SARS-CoV-2 viral attachment and infectivity is higher at low temperatures. (**A**) Pseudoviruses encoding the luciferase gene (Luc+) and bearing SARS-CoV-2 Spike (D614G or D614G N501Y) were tested for virus capture by ACE2-Fc at 37°C or 4°C. RLU obtained using ACE2-Fc were normalized to the signal obtained with the CV3-25 mAb. (**B**) Pseudoviruses Luc+ bearing SARS-CoV-2 Spike (D614G, D614G N501Y or B.1.1.7), or VSV-G as a control, were used to infect 293T-ACE2 cells. Virions were incubated at 37°C, 22°C or 4°C for 1h prior infection of 293T-ACE2 cells for 48h at 37°C. (**C**) Cell-to-cell fusion was measured between 293T effector cells expressing HIV-1 Tat and SARS-CoV-2 Spike (D614G or B.1.1.7) which were incubated at 37°C or 4°C for 1 h prior co-culture with TZM-bl-ACE2 target cells. (**B-C**) RLU were normalized to the signal obtained with cells pre-incubated at 37°C. (**D-E**) Pseudoviruses Luc+ bearing SARS-CoV-2 Spike (WT, D614G or B.1.1.7) were used to infect 293T-ACE2 cells in presence of increasing concentrations of sACE2 at 37°C for 1h prior infection of 293T-ACE2 cells. Fitted curves and IC_50_ values were determined using a normalized non-linear regression. (**F**) Authentic SARS-CoV-2 D614G virus was used to infect reconstituted human airway epithelia. Viral attachment was performed at 37°C or 4° and cells were further cultured at 37°C for 96h. Viral titers (RNA copies/mL) were monitored at 24h and 96h post-infection using one-step qRT-PCR. Viral titer values were normalized to the signal obtained with virions adsorbed to the cells at 37°C. Error bars indicate means ± SEM. These results were obtained in at least 3 independent experiments. Statistical significance was tested using (**A,C,F**) an unpaired T test or (**B**) one-way ANOVA with a Holm-Sidak post-test (*p < 0.05; **p < 0.01; ***p < 0.001; ****p < 0.0001; ns, non significant).

### Low temperatures enhance the affinity of SARS-CoV-2 RBD for ACE2

We then evaluated whether the impact of temperature on ACE2 interaction could be recapitulated by the RBD alone. Isothermal titration calorimetry (ITC) was used to measure the binding of ACE2 to RBD at different temperatures ranging from 10°C to 35°C (Fig S2A). The binding of ACE2 to RBD WT at 25°C is characterized by a dissociation constant (*K_D_*) of 19 nM in a process that is associated with a favorable change in enthalpy of – 20 kcal/mol, which is partially compensated by an unfavorable entropy contribution of 9.5 kcal/mol (Fig 3A & S2B). The data obtained at different temperatures reveal a 3-fold increase in binding affinity from 43 nM at 35°C to 14 nM at 15°C. The observed effect of the temperature on *K_D_* is expected based on the known temperature dependence of Gibbs energy of binding, *ΔG(T)*, which allows calculation of the expected *K_D_* values at any temperature (Fig S2C). Furthermore, as expected, the binding affinity of sACE2 to RBD N501Y was at least 6-fold higher than to RBD WT (Fig 3A). The *K_D_* value for the N501Y mutant is 2.9 nM at 25°C and the respective values for the enthalpy and entropy contributions are -16.6 and 4.9 kcal/mol. The gain in affinity is the result of a loss in unfavorable entropy which is larger than and overcompensates the loss in favorable enthalpy. Compared to ACE2 binding to RBD WT, the increase in binding affinity upon a drop in temperature is larger for the N501Y mutant with the *K_D_* changing from 6.9 nM at 35°C to 1 nM at 15°C.

**Figure 3.**
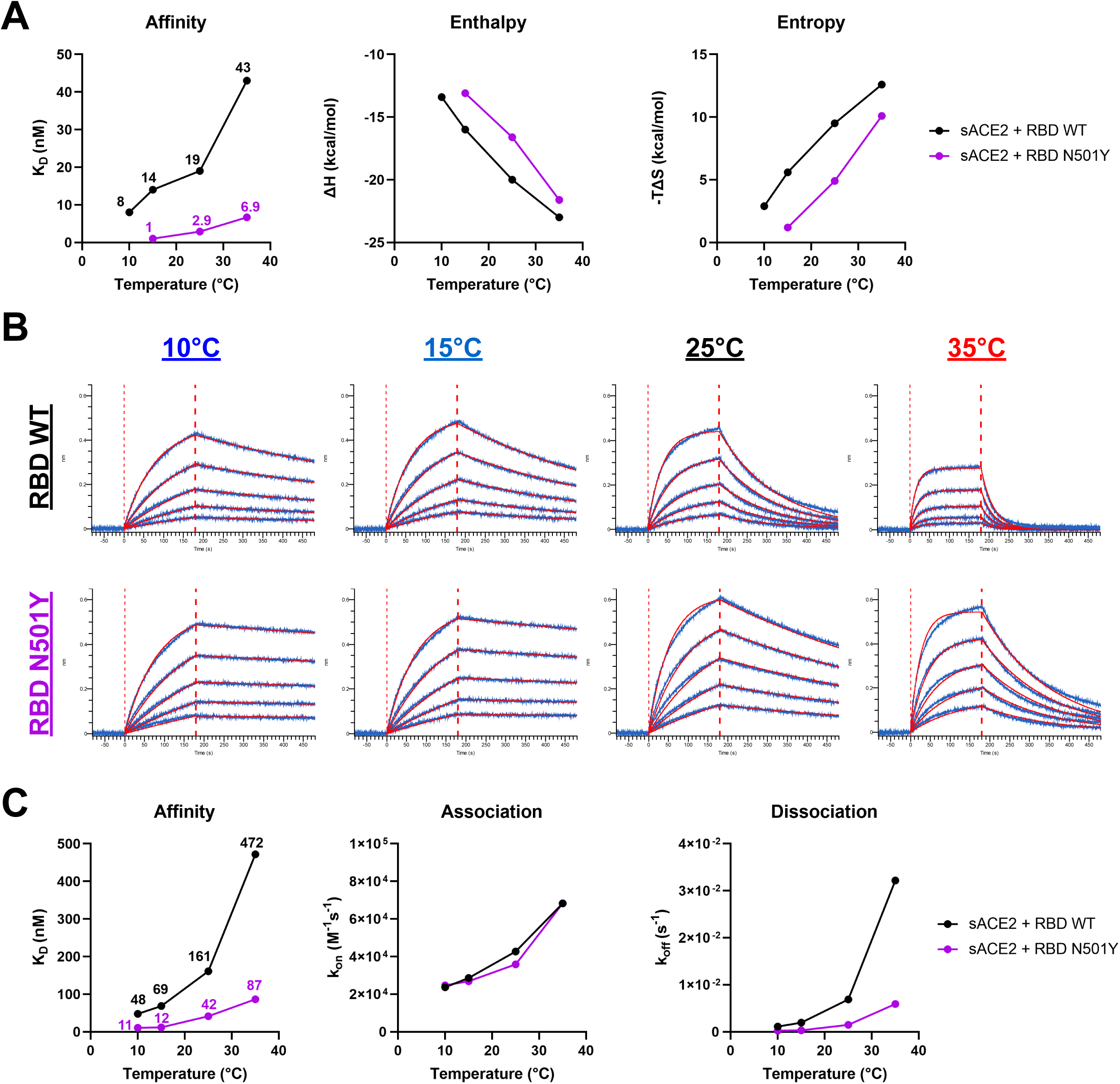
Enhanced affinity of SARS-CoV-2 RBD for ACE2 at low temperatures. (**A**) The thermodynamic parameters of sACE2 binding to SARS-CoV-2 RBD WT or N501Y measured by ITC at 10°C, 15°C, 25°C and 35°C. The graphs shown represent the affinity (*K_D_*), enthalpy (*ΔH*) and entropy (*-TΔS*) values obtained at the different temperatures. All ITC titration curves and thermodynamics values are shown in Figure S2. (**B-C**) Binding kinetics between SARS-CoV-2 RBD (WT or N501Y) and sACE2 assessed by BLI at different temperatures. (**B**) Biosensors loaded with RBD proteins were soaked in two-fold dilution series of sACE2 (500 nM to 31.25 nM) at different temperatures (10°C, 15°C, 25°C or 35°C). Raw data are shown in blue and fitting model is shown in red (**C**) Graphs represent the affinity constants (*K_D_*), on rates (*K_on_*) and off rates (*K_off_*) values obtained at different temperatures and calculated using a 1:1 binding model. All BLI data are summarized in Table S1.

To better characterize how the temperature affects the binding kinetics between RBD and ACE2, we used biolayer interferometry (BLI) at the same temperatures as for ITC. RBD proteins were immobilized on biosensors and were soaked in increasing concentrations of sACE2, ranging from 31.25 to 500 nM (Fig 3B). Again, affinity between RBD WT and sACE2 was found to be higher at lower temperatures (∼10-fold increase between 35°C and 10°C). Changes in affinity were explained by a major decrease in the off rate kinetics at low temperatures, despite a concomitant decrease in on rate kinetics (Fig 3C, Table S1). Compared to its WT counterpart, introduction of the N501Y mutation significantly decreased the off rate resulting in a 4.6 fold increase in *K_D_* when performed at 25°C. Remarkably, RBD WT reached a similar affinity for sACE2 at 10°C than the one achieved by RBD N501Y at 25°C (Fig 3B). Altogether, this indicates that low temperatures or the N501Y mutation confer analogous affinity changes that are favorable for Spike RBD-ACE2 interaction.

### Low temperatures modulate SARS-CoV-2 Spike trimer opening

Since ACE2 interaction with Spike occurs when its RBD is in the “up” conformation (50–52), we sought to determine if temperature could also be modulating Spike trimer opening (i.e. RBD accessibility). To do so, we evaluated the degree of cooperativity between sACE2 monomer binding within the Spike trimers by calculating the Hill coefficient (h), since ACE2 is thought to interact with Spike RBDs in a sequential manner (51). The h values are calculated from the steepness of dose-response curves generated upon incubation of Spike-expressing cells with increasing concentrations of sACE2 as previously described (43). We observed that the binding cooperativity of ACE2 to Spike D614G was slightly negative at 37°C (h=0.816), while being neutral at 4°C (h=1.004) (Fig 4A). On the contrary, the binding cooperativity to Spike B.1.1.7 was already slightly positive at 37°C (h=1.183) and was further improved at 4°C (h=1.371), suggesting that B.1.1.7 mutations could facilitate a coordinated Spike opening in addition to its increased ACE2-RBD interaction, thus fueling the viral entry process (Fig 4B). Spike conformational changes induced by temperature variation were also investigated by measuring the binding of the CR3022 mAb which specifically recognizes the RBD “up” conformation (53, 54). Despite no clear change in binding affinity to RBD at low temperatures, CR3022 bound better to the membrane-bound trimeric Spike at 4°C compared to 37°C (Fig 4C-D, Table S1). Since CR3022 is known to disrupt prefusion Spike trimer (RBD) (53, 54), we also confirmed this phenotype using an uncleaved Spike version (Furin KO) (Fig 4C). However, the increase in binding by CR3022 at 4°C was minor compared to the one observed with ACE2-Fc and no change was seen at 22°C whereas ACE2-Fc binding was significantly higher (Fig 1D & 4C). This confirms that low temperatures facilitate the exposure of the RBD in the “up” conformation, but it is unlikely sufficient on its own to recapitulate the temperature-dependent modulation of ACE2 interaction described in Figures 1 and 2.

**Figure 4.**
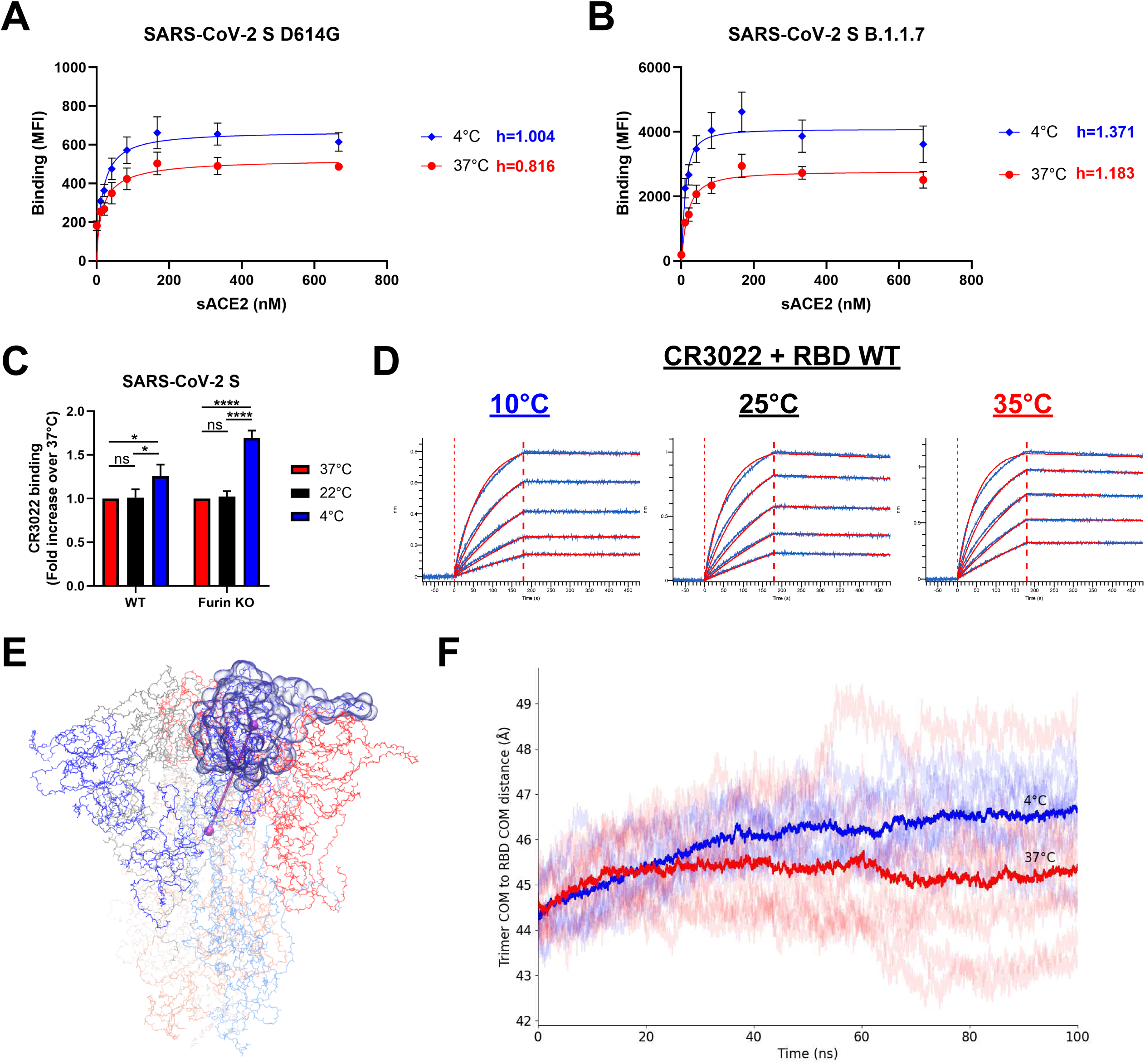
SARS-CoV-2 Spike trimer “opens” at low temperatures. (**A-B**) Binding of sACE2 to SARS-CoV-2 Spike (**A**) D614G or (**B**) B.1.1.7 expressed on 293T cells was measured at 37°C or 4°C by flow cytometry. Cells were preincubated with increasing amounts of sACE2 and its binding was detected using an anti-ACE2 staining. The Hill coefficients were determined using GraphPad software. (**C**) Cell-surface staining of transfected 293T cells expressing SARS-CoV-2 Spike (WT or Furin KO) using the CR3022 mAb when performed at 37°C, 22°C or 4°C. (**A-C**) The graphs shown represent the median fluorescence intensities (MFI). Error bars indicate means ± SEM. These results were obtained in at least 3 independent experiments. Statistical significance was tested using (**C**) one-way ANOVA with a Holm-Sidak post-test (*p < 0.05; ****p < 0.0001; ns, non significant). (**D**) Binding kinetics between RBD WT and CR3022 mAb assessed by BLI at 10°C, 25°C or 35°C. Biosensors loaded with RBD were soaked in two-fold dilution series of CR3022 (100 nM to 6.25 nM). Raw data are shown in blue and fitting model (1:1 binding model) is shown in red. All BLI data are summarized in Table S1. (**E)** Snapshot of SARS-CoV-2 Spike ectodomain (PDB 6VXX) (14) with one RBD indicated in transparent surface and one protomer’s RBD-to-trimer-center-of-mass distance indicated with a cylinder. **(F**) Traces of the RBD-to-trimer distances from three replicas each of all-atom, fully glycosylated and solvated MD simulations of the closed SARS-CoV-2 S trimer at 4°C (blues) and 37°C (reds) with dataset averages shown in heavy traces.

To better understand how low temperature affects the conformational dynamics of Spike and the propensity of RBD to sample the “up” conformation, we performed all-atom molecular dynamics (MD) simulations to measure the distance between the center of mass of the trimer and the center of mass of each RBD subunit using the structure of a fully glycosylated closed SARS-CoV-2 Spike ectodomain trimer as a model (Fig 4E) (14). Shown in Figure 4F is the RBD-to-trimer-center distances of all S1 subunits in three replicas for each temperature (37°C or 4°C). At 4°C, this distance is on average about 1.5 Å longer than at 37°C, suggesting that lower temperatures favor conformations that are, on average, closer to RBD opening than do higher temperatures. This quaternary structural sensitivity to temperature is consistent with the observation that CR3022 is more reactive against full Spike trimers, but not RBD alone, at lower temperatures (Fig 4C-F).

## DISCUSSION

In this study, we analyzed the role of temperature in modulating the affinity of SARS-CoV-2 Spike glycoprotein for its host receptor ACE2. We observed a significant enhancement in the affinity at low temperatures which could be explained by favorable thermodynamics changes leading to a stabilization of the RBD-ACE2 interface and by the triggering of more “open” conformations of the Spike trimer. Consequently, SARS-CoV-2 entry events and early replication kinetics were found to be amplified by enhanced viral adsorption at cold temperatures. This could potentially lead to higher transmissibility and faster replication in upper airway tissues upon exposure to the virus at lower seasonal temperatures. While we did not explore this possibility, temperature could also be affecting viral replication kinetics post-exposure and one could speculate that elevated body temperature resulting from SARS-CoV-2 infection (>38°C) could participate in limiting virus replication *in vivo* by interfering with viral entry, as previously suggested (55).

In summary, our results suggest that the RBD from the original strain isolated in Wuhan requires lower temperature for optimal interaction with ACE2 whereas the N501Y mutation frees RBD from this requirement. Whether this mechanism contributes to viral transmission and the apparent lack of seasonality for VOCs transmitted at warmer temperatures remains to be demonstrated. Our results indicate that the RBD-ACE2 affinity should be taken into consideration when evaluating the impact of the temperature on SARS-CoV-2 transmission.

## EXPERIMENTAL PROCEDURES

Experimental procedures are provided as supporting information.

## DATA AVAILABILITY

All data are contained within the article.

## SUPPORTING INFORMATION

This article contains supporting information (8,13,14,26,43–45,49,56–65).

## ACKNOWLEDGMENTS

The authors thank the CRCHUM BSL3 and Flow Cytometry Platforms for technical assistance. We thank Dr. Stefan Pöhlmann and Dr. Markus Hoffmann (Georg-August University) for the plasmids coding for SARS-CoV-2 and SARS-CoV-1 S, Dr. M. Gordon Joyce (U.S. MHRP) for the CR3022 mAb and Dr Jason McLellan for the plasmid coding for sACE2. The CV3-25 mAb was produced using the pTT vector kindly provided by the Canada Research Council.

## AUTHOR CONTRIBUTIONS

J.P. and A.F. conceived the study. J.P., J.R., C.F.A., A.S. and A.F. designed experimental approaches; J.P., J.R., R.G., S.D., C.F., S.P.A., A.T., M.B., N.G.V., G.B., C.F.A., A.S. and A.F. performed, analyzed, and interpreted the experiments; J.P., D.A., S.Y.G., G.G., A.P., S.M., H.C., M.R., W.M., M.P., E.B. and A.F. contributed unique reagents; J.P., C.F.A., A.S. and A.F. wrote the paper. Every author has read, edited, and approved the final manuscript.

## FUNDING AND ADDITIONAL INFORMATION

This work was supported by le Ministère de l’Économie et de l’Innovation (MEI) du Québec, Programme de soutien aux organismes de recherche et d’innovation to A.F., the Fondation du CHUM, a CIHR foundation grant #352417, an Exceptional Fund COVID-19 from the Canada Foundation for Innovation (CFI) #41027, the Sentinelle COVID Quebec network led by the Laboratoire de Santé Publique du Quebec (LSPQ) in collaboration with Fonds de Recherche du Québec-Santé (FRQS) and Genome Canada – Génome Québec, and by the Ministère de la Santé et des Services Sociaux (MSSS) and MEI. A.F. is the recipient of Canada Research Chair on Retroviral Entry no. RCHS0235 950-232424. J.P. and S.P.A. are supported by CIHR fellowships. R.G. is supported by a MITACS Accélération postdoctoral fellowship. This work used the Extreme Science and Engineering Discovery Environment (XSEDE) (66) resource Stampede2 at the Texas Advanced Computing Center through allocation TG-MCB070073N. XSEDE is supported by National Science Foundation grant number ACI-1548562. G.B. is the recipient of a CIHR foundation grant. The funders had no role in study design, data collection and analysis, decision to publish, or preparation of the manuscript.

## DISCLAIMER

The views expressed in this presentation are those of the authors and do not reflect the official policy or position of the Uniformed Services University, US Army, the Department of Defense, or the US Government.

## SUPPLEMENTAL MATERIAL

### EXPERIMENTAL PROCEDURES

#### Ethics statement

Primary airway epithelial cells (AECs), isolated from lung biopsies collected from healthy individuals were provided by the CRCHUM’s Respiratory Cell and Tissue Biobank from the Respiratory Health Research Network of Québec with informed written consent prior to enrollment in accordance with Institutional Review Board approval (#CE08.063) and approval of the research study by the CRCHUM institutional review board (protocol #20.454). Research adhered to the standards indicated by the Declaration of Helsinki.

#### Sequence alignment

The Logo plots (1) were created using the WebLogo program (https://weblogo.berkeley.edu/logo.cgi) and the COVID-19 CoV Genetics interface (2) (https://covidcg.org/) using the GISAID database (https://www.gisaid.org/) to identify single-nucleotide polymorphism (SNP). The alignments correspond to the worldwide available sequences deposited between December 15^th^, 2019 and December 31^st^, 2020 (2019-2020 alignment) or January 1^st^, 2021 and June 18^st^, 2021 (2021 alignment), which includes 486,432 and 1,195,746 individual sequences of SARS-CoV-2 Spike RBD (residues 319-541), respectively. The relative height of each letter within individual stack represents the frequency of the indicated amino acid at that position. The numbering of all the Spike amino acid sequences is based on the prototypic WIV04 strain of SARS-CoV-2, where 1 is the initial methionine (3).

#### Cell lines, primary cells and viruses

293T human embryonic kidney cells (ATCC), Vero E6 african green monkey kidney cells (ATCC), Cf2Th (ATCC), 293T-ACE2 and TZM-bl-ACE2 cells were maintained at 37°C under 5% CO_2_ in Dulbecco’s Modified Eagle Medium (DMEM) (Wisent), supplemented with 5% fetal bovine serum (FBS) (VWR) and 100 U/mL penicillin/streptomycin (Wisent). 293T-ACE2 and TZM-bl-ACE2 cells stably expressing human ACE2 are derived from 293T cells and TZM-bl cells, respectively (4, 5). Cf2Th cells are SARS-CoV-2-resistant canine thymocytes and were used in the virus capture assay. 293T-ACE2 and TZM-bl-ACE2 cells were then cultured in medium supplemented with 2 µg/mL of puromycin (Millipore Sigma). Primary human airway epithelial cells (AECs) used in Figure 1F were isolated from bronchial biopsies collected from two healthy subjects (two males, mean age of 49 yrs). Briefly, bronchial tissues were rinsed and then incubated overnight at 4°C with MEM medium (Life Technologies) supplemented with 7.5% NaHCO_3_ (Sigma-Aldrich), 2 mM L-glutamine, 10 mM HEPES (ThermoFisher Scientific), 0.05 mg/ml gentamycin, 50 U/ml penicillin-streptomycin, 0.25 μg/ml Fungizone (Life Technologies) and containing 0.1% protease (from Streptomyces griseus; Sigma-Aldrich) and 10 μg/ml DNAse (Deoxyribonuclease I from bovine pancreas; Sigma-Aldrich). The protease and DNAse activities were then neutralized with FBS (Life Technologies). AECs were gently scraped off the remaining tissue and red blood cells were removed by treatment with ACK lysis buffer (0.1 mM NH_4_Cl, 10 μM KHCO_3_, 10 nM EDTA). After counting, the freshly isolated cells were seeded into flasks coated with Purecol (Cedarlane) and collagen IV (Sigma-Aldrich), and grown in a mix (50:50) of PneumacultEx (STEMCELL Technologies) and CnT-17 (CellnTec Advanced Cell Systems) media for two days, and then in CnT-17 until the confluence is reached.

AECs were then detached with a trypsin solution before seeding into 6 wells plates coated with Purecol and collagen IV, cultured in CnT-17 until confluency (∼5 days) ad then in BEGM (Lonza) for two days before experimentations. Authentic SARS-CoV-2 viruses used in Figure 1F were isolated, sequenced and amplified from clinical samples obtained from patients infected with SARS-CoV-2 D614G or B.1.1.7 by the Laboratoire de Santé Publique du Québec (LSPQ). SARS-CoV-2/Québec/CHUL/21697 is a clinical sample isolated in Quebec City, Canada and then amplified on Vero E6 cells. Virus was sequenced by MinION technology (Oxford Nanopore technologies, Oxford, UK). All work with infectious using SARS-CoV-2 authentic virus was performed in a Biosafety Level 3 (BSL3) facilities at Université de Montréal and Université Laval using appropriate positive pressure air respirators and protective equipments.

#### Plasmids and site-directed mutagenesis

The plasmids expressing the wildtype SARS-CoV-2 and SARS-CoV-1 Spikes were previously reported (6, 7). The plasmid encoding for SARS-CoV-2 S RBD (residues 319-541) fused with a hexahistidine tag was previously described (8). The mutations in full-length Spike (D614G, N501Y and/or R682S/R683S) or RBD (N501Y) expressors were introduced using the QuikChange II XL site-directed mutagenesis protocol (Agilent Technologies). The presence of the desired mutations was determined by automated DNA sequencing. The plasmid encoding the SARS-CoV-2 Spike from the B.1.1.7 lineage (Δ69-70, Δ144, N501Y, A570D, D614G, P681H, T716I, S982A and D1118H) was codon-optimized and synthesized by Genscript (9). The plasmid encoding for soluble human ACE2 (residues 1–615) fused with an 8xHisTag was reported elsewhere (10). The plasmid encoding for the ACE2-Fc chimeric protein, a protein composed of an ACE2 ectodomain (1–615) linked to an Fc segment of human IgG1 was previously reported (11). The vesicular stomatitis virus G (VSV-G)-encoding plasmid (pSVCMV-IN-VSV-G) was previously described (12).

#### Protein expression and purification

FreeStyle 293F cells (Thermo Fisher Scientific) were grown in FreeStyle 293F medium (Thermo Fisher Scientific) to a density of 1 x 10^6^ cells/mL at 37°C with 8 % CO_2_ with regular agitation (150 rpm). Cells were transfected with a plasmid coding for SARS-CoV-2 S RBD, sACE2 or ACE2-Fc using ExpiFectamine 293 transfection reagent, as directed by the manufacturer (Thermo Fisher Scientific). One week later, cells were pelleted and discarded. Supernatants were filtered using a 0.22 µm filter (Thermo Fisher Scientific). The recombinant RBD and sACE2 proteins were purified by nickel affinity columns, as directed by the manufacturer (Invitrogen). The ACE2-Fc chimeric protein was purified by protein A affinity columns, as directed by the manufacturer (Cytiva). The recombinant protein preparations were dialyzed against phosphate-buffered saline (PBS) and stored in aliquots at -80°C until further use. To assess purity, recombinant proteins were loaded on SDS-PAGE gels and stained with Coomassie Blue.

#### Flow cytometry analysis of cell-surface staining (transfected cells)

Using the standard calcium phosphate method, 10 μg of Spike expressor and 2 μg of a green fluorescent protein (GFP) expressor (pIRES2-eGFP; Clontech) was transfected into 2 × 10^6^ 293T cells. At 48h post transfection, 293T cells were stained with anti-Spike monoclonal antibodies CV3-25 (13) or CR3022 (14) (5 µg/mL) or using the ACE2-Fc chimeric protein (20 µg/mL) for 45 min at 37℃, 22℃ or 4℃. Alternatively, to determine the Hill coefficients, cells were preincubated with increasing concentrations of sACE2 (0 to 719 nM) at 37°C or 4°C. sACE2 binding was detected using a polyclonal goat anti-ACE2 (RND systems). Alexa Fluor-647- conjugated goat anti-human IgG (H+L) Abs (Invitrogen) and Alexa Fluor-647-conjugated donkey anti-goat IgG (H+L) Ab (Invitrogen) were used as secondary antibodies to stain cells for 30 min at room temperature. The percentage of transfected cells (GFP+ cells) was determined by gating the living cell population based on the basis of viability dye staining (Aqua Vivid, Invitrogen). Samples were acquired on a LSRII cytometer (BD Biosciences) and data analysis was performed using FlowJo v10.5.3 (Tree Star). Hill coefficient analyses were done using GraphPad Prism version 9.1.0 (GraphPad).

#### Flow cytometry analysis of cell-surface staining (infected cells)

SARS-CoV-2 authentic viruses (D614G or B.1.1.7 variant) were used to infect Vero E6 cells or primary AECs at a multiplicity of infection (MOI) of 0.0001 or 0.1, respectively. At 48h post-infection, cells were detached by PBS-EDTA and were stained with Abs for 30 min at 37℃ or 4℃. Alexa Fluor-647-conjugated goat anti-human IgG (H+L) Ab (Invitrogen) was used as secondary antibody to stain cells for 30 min at room temperature. Cells were then fixed with PBS containing 4% paraformaldehyde for 48h at 4℃. Then the cells were stained intracellularly for SARS-CoV-2 nucleocapsid (N) antigen, using the Cytofix/Cytoperm fixation/permeabilization kit (BD Biosciences) and an anti-N mAb (clone mBG17; Kerafast) conjugated with the Alexa Fluor 488 dye according to the manufacturer’s instructions (Invitrogen). The percentage of infected cells (N+ cells) was determined by gating the living cell population based on the basis of viability dye staining (Aqua Vivid, Invitrogen). Samples were acquired on a LSRII cytometer (BD Biosciences) and data analysis was performed using FlowJo v10.5.3 (Tree Star).

#### Virus capture assay

The SARS-CoV-2 virus capture assay was previously reported (15). Briefly, pseudoviral particles were produced by transfecting 2×10^6^ 293T cells with pNL4.3 Luc R-E- (3.5 μg), plasmids encoding for SARS-CoV-2 Spike (3.5 μg) proteins and VSV-G (1 μg) using the standard calcium phosphate method. Forty-eight hours later, virus-containing supernatants were collected, and cell debris were removed through centrifugation (1,500 rpm for 10 min). CV3-25 antibodies or ACE2-Fc proteins were immobilized on white MaxiSorp ELISA plates (Thermo Fisher Scientific) at a concentration of 5 μg/mL in 100 μL of PBS overnight at 4°C. Unbound proteins were removed by washing the plates twice with PBS. Plates were subsequently blocked with 3% bovine serum albumin (BSA) in PBS for 1 hour at room temperature, followed by 1 hour incubation at 37℃ or 4℃. Meanwhile, virus-containing supernatants were pre-tempered at 37°C or 4°C for 1 hour. After washing plates twice with PBS, 200 μL of virus-containing supernatant were added to the wells. After 30 min of incubation at 37°C or 4°C, supernatants were discarded, and the wells were washed with PBS three times. Virus capture was visualized by adding 1×10^4^ SARS-CoV-2-resistant Cf2Th cells per well in complete DMEM. Forty-eight hours post-infection, cells were lysed by the addition of 30 μL of passive lysis buffer (Promega) and one freeze-thaw cycle. An LB942 TriStar luminometer (Berthold Technologies) was used to measure the luciferase activity of each well after the addition of 100 μL of luciferin buffer (15 mM MgSO_4_, 15 mM KH_2_PO_4_ [pH 7.8], 1 mM ATP, and 1 mM dithiothreitol) and 50 μL of 1 mM D-luciferin potassium salt (ThermoFisher Scientific).

#### Pseudovirus infectivity and neutralization assay

293T-ACE2 target cells were infected with single-round luciferase-expressing lentiviral particles (4). Briefly, 293T cells were transfected by the calcium phosphate method with the lentiviral vector pNL4.3 R-E-Luc (NIH AIDS Reagent Program) and a plasmid encoding for SARS-CoV-2 Spike or VSV-G at a ratio of 5:4. Two days post-transfection, cell supernatants were harvested and used fresh for infectivity measurements or stored at –80°C until use for virus neutralization measurements. 293T-ACE2 target cells were seeded at a density of 1×10^4^ cells/well in 96-well luminometer-compatible tissue culture plates (Perkin Elmer) 24h before infection. To assess pseudovirus infectivity, freshly produced recombinant viruses were incubated for 1h at 37°C, 22°C or 4°C and were added to the target cells followed by incubation for 48h at 37°C. To measure virus neutralization by sACE2, recombinant viruses in a final volume of 100 μl were incubated with the increasing sACE2 concentrations (0 to 12,000 nM) for 1h at 37°C or 4°C and were then added to the target cells followed by incubation for 48h at 37°C; cells were lysed by the addition of 30 μl of passive lysis buffer (Promega) followed by one freeze-thaw cycle. An LB942 TriStar luminometer (Berthold Technologies) was used to measure the luciferase activity of each well after the addition of 100 μl of luciferin buffer (15 mM MgSO_4_, 15 mM KH_2_PO_4_ [pH 7.8], 1 mM ATP, and 1 mM dithiothreitol) and 50 μl of 1 mM D-luciferin potassium salt (ThermoFisher Scientific). The neutralization half-maximal inhibitory dilution (IC_50_) represents the sACE2 concentration inhibiting 50% of the infection of 293T-ACE2 cells by recombinant viruses bearing the indicated surface glycoproteins at different temperatures.

#### Cell-to-cell fusion assay

To assess the cell-to-cell fusion between Spike-expressing effector cells and ACE2-expressing target cells (5), 2×10^6^ 293T cells were co-transfected with plasmid expressing HIV-1 Tat (1 µg) and a plasmid expressing SARS-CoV-2 Spike (4 µg) using the calcium phosphate method. Two days after transfection, Spike-expressing 293T (effector cells) were detached with PBS-EDTA 1 mM and incubated for 1 h at 37°C or 4°C. Subsequently, effector cells (1 × 10^4^) were added to TZM-bl-ACE2 target cells that were seeded at a density of 1×10^4^ cells/well in 96-well luminometer-compatible tissue culture plates 24 h before the assay. Cells were co-incubated for 6 h at 37°C and 5% CO_2_, after which they were lysed by the addition of 40 μl of passive lysis buffer (Promega) and one freeze thaw cycle. An LB942 TriStar luminometer (Berthold Technologies) was used to measure the luciferase activity of each well after the addition of 100 μl of luciferin buffer (15 mM MgSO_4_, 15 mM KH_2_PO_4_ [pH 7.8], 1 mM ATP, and 1 mM dithiothreitol) and 50 μL of 1 mM D-luciferin potassium salt (ThermoFisher Scientific).

#### Viral infection of reconstituted human airway epithelia (MucilAir)

For this experiment, an *ex vivo* system was used. This consists of an air liquid interface that mimicks the human upper airway epithelium. The primary nasal cells were isolated from a pool of 14 donors (MucilAir, Epithelix). The experiment was performed in quadruplicate and cells were cultured in medium provided by manufacturer. Before infection, MucilAir wells were washed with warm OptiMEM medium (ThermoFisher Scientific) and then were pre-incubated at 4°C or 37°C for 15 min. Apical poles were then infected directly with 200 μL of virus (SARS-CoV-2/Québec/CHUL/21697) at a multiplicity of infection (MOI) of 0.015 and then incubated at 4°C or 37°C during 30 min. After this adsorption phase, virus-containing medium was removed and all MucilAir wells were placed at 37°C. Samples were collected from apical washes (200 μL of OptiMEM) at different timepoints post-infection (24h and 96h) and 100 μL were used to extract RNA (MagNA Pure LC, Total nucleic acid isolation kit, Roche Applied Science). Viral titers were determined by RT-qPCR one-step (QuantiTect Virus +ROX Vial Kit, Qiagen). To monitor the viability and health conditions of infected primary epithelial cells, transepithelial electrical resistance (TEER) was measured using a dedicated volt-ohm meter (Millicell® ERS-2, Millipore Sigma) and no significant difference was observed between cells infected at 4°C or 37°C.

#### Isothermal Titration Calorimetry (ITC)

Calorimetric titration experiments were performed using a MicroCal VP-ITC (Malvern Panalytical). The reagents were prepared in PBS pH 7.4, and then exhaustively dialyzed prior to the experiments. Any further dilutions of the reagents were made using the dialysate to avoid any unnecessary heats of dilution associated with the injections. All reagents were thoroughly degassed prior to the experiments. The enthalpy and affinity of binding at each temperature were determined from a complete titration of either RBD WT or the N501Y mutant with sACE2. The sACE2 solution at 11 - 13 µM was injected in 10 µL aliquots into the stirred calorimetric cell (v ∼ 1.4 mL) containing RBD protein at ∼ 1.5 µM. The titrations of RBD WT were performed at 10°C, 15°C, 25°C, and 35°C, while the titrations of the N501Y mutant were performed at 15°C, 25°C, and 35°C. The injection peaks were integrated, and the heat associated with binding was obtained after subtraction of the heats of dilution. The association constant, *K_A_* (the dissociation constant, *K_D_* =1/*K_A_*), the enthalpy change, *ΔH*, and the stoichiometry, *N*, were obtained by nonlinear regression of the data to a single-site binding model using Origin with a fitting function made in-house. Gibbs energy, *ΔG*, was calculated from the binding affinity using *ΔG* = -*RT*ln*K_A_*, (*R* = 1.987 cal/(K × mol)) and *T* is the absolute temperature in kelvin). The entropy contribution to Gibbs energy, *-TΔS*, was calculated from the relation *ΔG = ΔH -TΔS*. The Gibbs energy of binding, *ΔG(T)*, as a known temperature dependence which allows calculation of the *K_D_* values at any temperature according to:

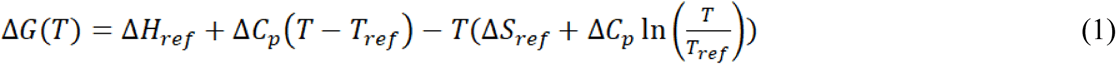

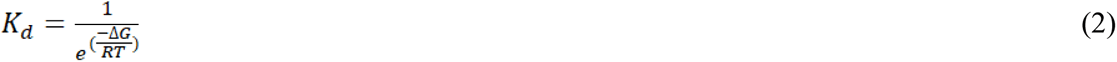

where *ΔH_ref_* and *ΔS_ref_* are the respective enthalpy and entropy changes at the known reference temperature *T_ref_*, *T* is the absolute temperature in kelvin, *R* the gas constant (1.987 cal/(K × mol)), and *ΔC_p_* the change in heat capacity upon binding. The *ΔC_p_* value for ACE2 binding is - 380 cal/(K × mol) for RBD WT and -425 cal/(K × mol) for RBD N501Y, which is obtained from the temperature dependence of the enthalpy change (Fig 3A). Figure S2C shows a plot of the experimental dissociation constants at different temperatures together with the expected values calculated from *ΔC_p_* and the enthalpy change at 25°C.

#### Biolayer interferometry (BLI)

Binding kinetics were performed on using an Octet RED96e system (ForteBio) at different temperatures (10°C, 15°C, 25°C, 35°C) with shaking at 1,000 RPM. Amine Reactive Second-Generation (AR2G) biosensors were hydrated in water, then activated for 300 s with a solution of 5 mM sulfo-NHS and 10 mM EDC (ForteBio) prior to amine coupling. Either SARS-CoV-2 RBD WT or the N501Y mutant were loaded into AR2G biosensor at 12.5 µg/mL at 25°C in 10 mM acetate solution pH 5 (Fortebio) for 600 s then quenched into 1 M ethanolamine solution pH 8.5 (Fortebio) for 300 s. Loaded biosensor were placed in 10X kinetics buffer (ForteBio) for 120 s for baseline equilibration. Association of sACE2 (in 10X kinetics buffer) to the different RBD proteins was carried out for 180 s at various concentrations in a two-fold dilution series from 500 nM to 31.25 nM prior to dissociation for 300 s. The data were baseline subtracted prior to fitting performed using a 1:1 binding model and the ForteBio data analysis software. Calculation of on rates (k_on_), off rates (k_off_), and affinity constants (K_D_) was computed using a global fit applied to all data. Alternatively, affinity of the CR3022 mAb at various concentrations in a two-fold dilution series from 100 nM to 6.25 nM for the immobilized SARS-CoV-2 RBD WT was assessed at different temperatures (10°C, 25°C, 35°C).

#### Molecular dynamics simulations

A fully glycosylated model of the closed SARS-CoV-2 S ectodomain trimer was built from the 6VXX PDB entry (16). Six independent replicas with explicit waters were generated. Three were run for 100 ns at 37°C using NAMD 2.14 (17) (2 fs time step, Langevin thermostat with 5/ps frequency) and three were run for 100 ns at 4°C. The CHARMM36 force-field (18) and TIP3P water model were used. Configuration snapshots were saved every 2 ps. The instantaneous distance between the center of mass of the trimer and the center of mass of each receptor binding domain (residues 330 to 521 in each S1 subunit) was measured.

#### Statistical Analysis

Statistics were analyzed using GraphPad Prism version 9.1.0 (GraphPad, San Diego, CA, USA). Every data set was tested for statistical normality and this information was used to apply the appropriate (parametric or nonparametric) statistical test. P values <0.05 were considered significant; significance values are indicated as * P<0.05, ** P<0.01, *** P<0.001, **** P<0.0001.

## SUPPLEMENTAL TABLE

**Table S1.**
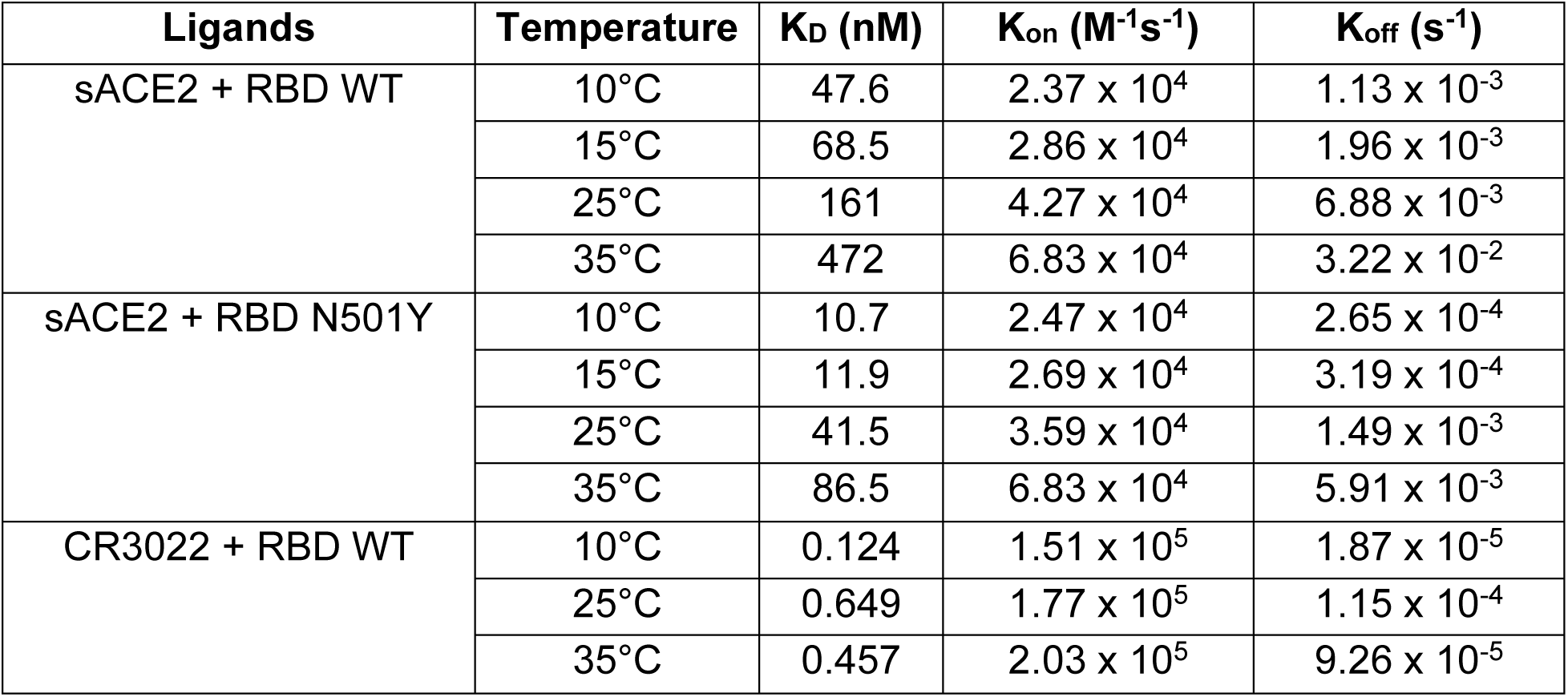
Affinity between SARS-CoV-2 RBD and its ligands quantified by biolayer interferometry.

| **Ligands** | **Temperature** | **K<sub>D</sub> (nM)** | **K<sub>on</sub> (M<sup>-1</sup>s<sup>-1</sup>)** | **K<sub>off</sub> (s<sup>-1</sup>)** |
| --- | --- | --- | --- | --- |
| sACE2 + RBD WT | 10°C | 47.6 | 2.37 x 10<sup>4</sup> | 1.13 x 10<sup>-3</sup> |
|  | 15°C | 68.5 | 2.86 x 10<sup>4</sup> | 1.96 x 10<sup>-3</sup> |
|  | 25°C | 161 | 4.27 x 10<sup>4</sup> | 6.88 x 10<sup>-3</sup> |
|  | 35°C | 472 | 6.83 x 10<sup>4</sup> | 3.22 x 10<sup>-2</sup> |
| sACE2 + RBD N501Y | 10°C | 10.7 | 2.47 x 10<sup>4</sup> | 2.65 x 10<sup>-4</sup> |
|  | 15°C | 11.9 | 2.69 x 10<sup>4</sup> | 3.19 x 10<sup>-4</sup> |
|  | 25°C | 41.5 | 3.59 x 10<sup>4</sup> | 1.49 x 10<sup>-3</sup> |
|  | 35°C | 86.5 | 6.83 x 10<sup>4</sup> | 5.91 x 10<sup>-3</sup> |
| CR3022 + RBD WT | 10°C | 0.124 | 1.51 x 10<sup>5</sup> | 1.87 x 10<sup>-5</sup> |
|  | 25°C | 0.649 | 1.77 x 10<sup>5</sup> | 1.15 x 10<sup>-4</sup> |
|  | 35°C | 0.457 | 2.03 x 10<sup>5</sup> | 9.26 x 10<sup>-5</sup> |

## SUPPLEMENTAL FIGURE CAPTIONS

**Figure S1.**
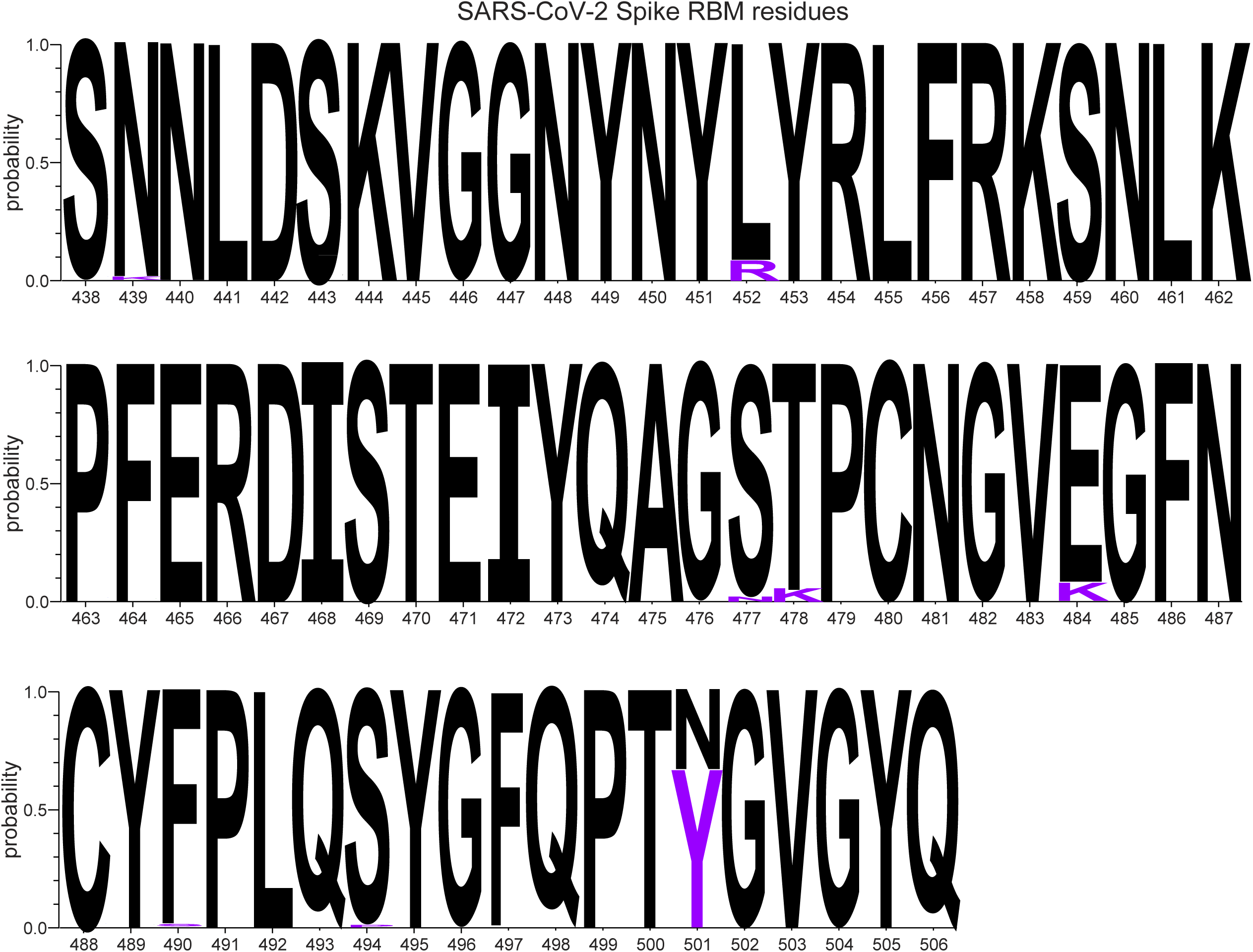
Sequence conservation of RBM residues among SARS-CoV-2 Spike isolates. Logo depictions of the frequency of selected SARS-CoV-2 Spike residues from the receptor binding motif (RBM). Worldwide sequences deposited in the GISAID database in 2021 (January 1^st^, 2021 to June 18^th^, 2021) were aligned using the COVID CoV Genetics program, which includes 1,195,746 individual sequences. Residue numbering is based on the SARS-CoV-2 WIV04 reference strain. The height of the letter indicates its frequency of total deposited sequences. Residues corresponding to the WIV04 reference sequence are shown in black and residues corresponding to emerging VOCs are shown in violet.

**Figure S2.**
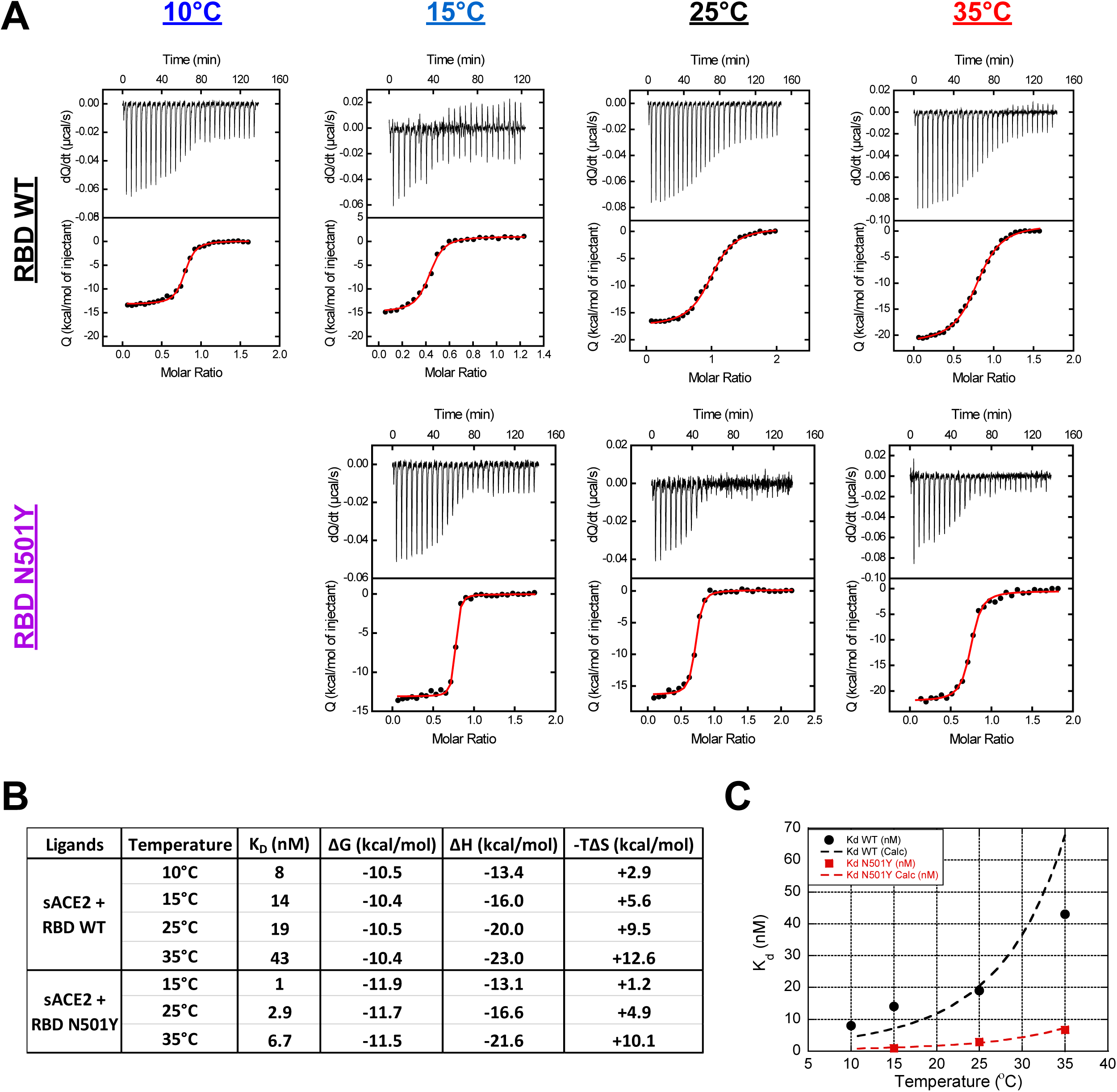
Titration of sACE2 binding to SARS-CoV-2 RBD by isothermal titration calorimetry. (**A**) ITC data from calorimetric titrations of SARS-CoV-2 RBD (WT and N501Y) with sACE2 at different temperatures. The top panel of each graph shows the heat flow, dQ/dt, as a function of time and the bottom panel shows the integrated heat associated with each injection (Q) as a function of the molar ratio between the concentrations of sACE2 and RBD in the cell. The solid line represents the result from best nonlinear least squares fit of the data to a single-site binding model. (**B**) Table summarizing the thermodynamic parameters for the binding of sACE2 to SARS-CoV-2 RBD (WT and N501Y). The values for the dissociation constant, *K_D_*, and enthalpy, *ΔH*, were obtained directly from nonlinear regression of the ITC data (**A**), while Gibbs energy, *ΔG*, and the entropy contribution, *-TΔS*, were calculated as described in the method section. (**C**) The *K_D_* values for the binding of sACE2 to SARS-CoV-2 RBD WT (filled circles) and N501Y (filled squares) as a function of temperature. The dashed lines correspond to the expected values calculated from *ΔC_p_* and the enthalpy change at 25 °C.

